# A genetically encoded fluorescent sensor for monitoring spatiotemporal prostaglandin E2 dynamics *in vivo*

**DOI:** 10.1101/2025.06.01.657218

**Authors:** Lei Wang, Yini Yang, Fei Deng, Yuqi Yan, Yulong Li

## Abstract

Prostaglandin E2 (PGE2) is an important lipid signaling molecule that regulates a wide range of physiological and pathological processes. However, its dynamics during these processes are largely unknown due to the lack of tools to directly visualize PGE2 with high spatiotemporal resolution. Here, we developed and characterized a genetically encoded PGE2 sensor, which we call GRAB_PGE2-1.0_ (PGE2-1.0), that has high specificity for PGE2, nanomolar affinity, rapid kinetics, and high spatial resolution when expressed both *in vitro* and *in vivo*. Using fiber-photometry recordings, we found that PGE2-1.0 can reliably monitor endogenous PGE2 dynamics in the preoptic area in the brain during acute inflammation. The wide-field *in vivo* imaging with PGE2-1.0 reveals spatial heterogeneity in cortex-wide PGE2 dynamics during acute inflammation and seizure. Thus, our PGE2-1.0 sensor can be used to detect endogenous PGE2 dynamics with high spatiotemporal resolution, providing a robust tool for studying PGE2 under specific physiological and pathological conditions.

## INTRODUCTION

Prostaglandins play an important role in regulating a variety of physiological and pathological processes. Prostaglandin E2 (PGE2)—the most abundant prostaglandin in the human body—is produced by a wide range of cell types in a variety of organisms^1,2^. PGE2 is closely associated with the onset and progression of inflammation, pain, fever, and neurological diseases; inhibiting prostaglandin production has long been an important strategy for improving anti-inflammatory and analgesic treatments^3,4^. PGE2 is synthesized from arachidonic acid by cyclooxygenase (COX) and prostaglandin E synthase (PTGES) enzyme families^5–7^, with COX enzymes serving as the key target for non-steroidal anti-inflammatory drugs (NSAIDs)^3^. Inhibiting COX activity— particularly COX-2 activity—to reduce PGE2 production is a widely used clinical strategy to alleviate pain, inflammation, and fever^3^.

PGE2 exerts its biological functions primarily by binding to and activating its four receptors (EP1, EP2, EP3, and EP4), which are widely expressed throughout the central nervous system. Importantly, however, the expression levels of these receptors differ across various brain regions^8–10^, and these receptors have different affinities for PGE2^11^. Thus, monitoring PGE2 dynamics will serve as an important step toward understanding the mechanism underlying PGE2’s various functions; however, the spatiotemporal dynamics of PGE2 remain largely unknown due to the lack of suitable tools for detecting PGE2 with high specificity, sensitivity, and spatiotemporal resolution. Unlike classic neurotransmitters stored in synaptic vesicles, evidences suggest that PGE2 is not stored within the cell but is synthesized and released on demand^12,13^. Hence, the expression levels of COX-2 and PTGES (particularly PTGES1) enzymes^14,15,5^ are widely used as an indirect measure of PGE2 levels, as methods used to directly detection PGE2 lack sufficient spatiotemporal resolution. For example, measuring PGE2 usually requires the extraction of tissue or blood samples, followed by purification and analysis steps, including radioimmunoassay^16^, enzyme-linked immunosorbent assay^17,18^, chromatography, and mass spectrometry; although this approach can provide precise information regarding PGE2 levels, it has poor spatiotemporal resolution and cannot be used for real-time recording *in vivo*. On the other hand, microdialysis can measure molecules in the brain with high specificity, but is difficult to use to detect hydrophobic molecules such as PGE2 and has low spatial and temporal resolution (with a sampling interval typically ranging from 20 to 30 min)^19,20^. Therefore, a genetically encoded tool to specifically detect PGE2 with high spatiotemporal resolution *in vivo* would fill a significant need in this field.

In recent years, many genetically encoded fluorescent sensors were developed to visualize specific neurochemicals using minimally invasive methods and providing high spatiotemporal resolution. One highly successful example of this approach is the GPCR activation-based (GRAB) strategy, in which a series of genetically encoded fluorescent sensors were developed based on various G protein-coupled receptors (GPCRs) and conformation-sensitive circularly permuted enhanced green fluorescent protein (cpEGFP), thus allowing the user to detect specific GPCR ligands such as neurotransmitters and neuromodulators with high specificity and high sensitivity^21–27^. Here, we used this strategy to develop a genetically encoded fluorescent sensor specific for PGE2. We found that this sensor, which we call GRAB_PGE2-1.0_ (hereafter PGE2-1.0), has high specificity for PGE2, high sensitivity (with a peak change in fluorescence >1000% in response to PGE2), rapid kinetics, and negligible coupling to downstream signaling. We then examined the *in vitro* performance of PGE2-1.0 expressed in cultured cells and acute mouse brain slices. Finally, we expressed PGE2-1.0 in the mouse brain and monitored extracellular PGE2 dynamics during both inflammation and seizure *in vivo*.

## RESULTS

### Development and characterization of a GRAB_PGE2_ sensor

To measure PGE2 dynamics with high spatial and temporal resolution, we developed a genetically encoded GRAB sensor for PGE2 (Figure 1A) based on the GPCR activation-based (GRAB) strategy. First, we systematically searched for the most suitable GPCR scaffold from four known human PGE2 receptors, namely the EP1, EP2, EP3, and EP4 receptors. For each GPCR, the third intracellular loop (ICL3) was replaced with the cpEGFP-containing ICL3 derived from our previously well-characterized green florescent norepinephrine sensor GRAB_NE1m_^23^, using various insertion sites. Candidate sensors were then screened for suitable membrane trafficking (Figure S1A–C) and change in fluorescence (ΔF/F_0_) in response to PGE2 (Figure S1D). After this initial screening, we identified a human EP2-based candidate as the prototype sensor, which we call PGE2-0.1 (Figure S1D). Next, to improve the sensor’s response, we conducted three optimization steps: *i*) fine tuning the ICL3 insertion sites, *ii*) truncating the linker region between the EP2 scaffold and cpEGFP, and *iii*) introducing mutations in the linker region and critical residues in cpEGFP. After screening more than 2000 variants, we identified PGE2-1.0 as the variant with the largest response (∼700%) to 1 μM PGE2 (Figures 1B and S2). In addition, based on the human EP2 structure^28^, we performed a mutagenesis screen of eight amino acids located near the receptor’s ligand-binding pocket and generated a PGE2-insensitive sensor (for use as a negative control) containing a single missense mutation (T113^2.54^W) in PGE2-1.0; we call this ligand-insensitive sensor PGE2mut (Figure 1B).

**Figure 1.**
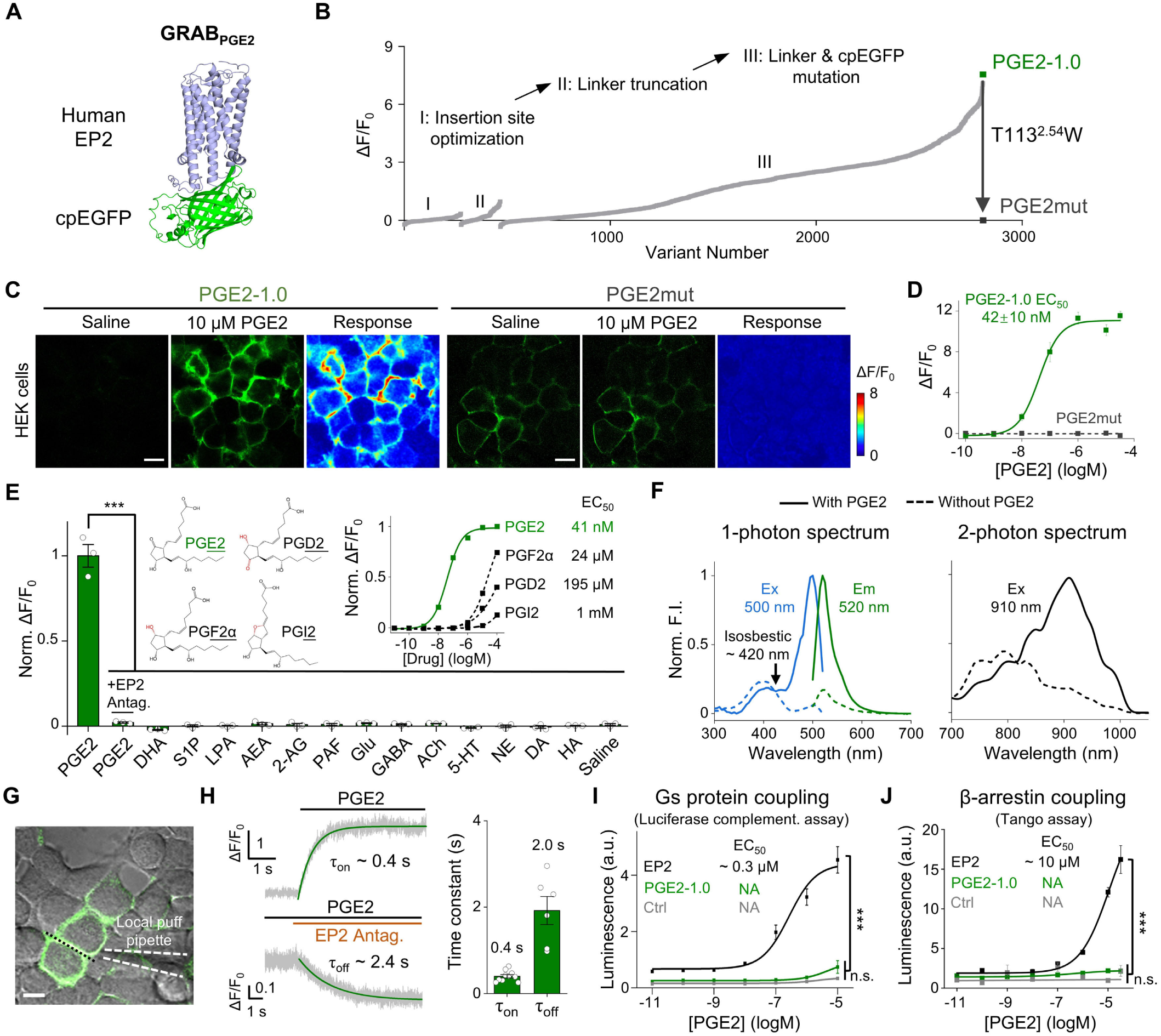
Development, optimization, and characterization of PGE2-1.0 in HEK293T cells. **(A)** Drawing depicting the structure of the GRAB_PGE2_ sensor, consisting of the human EP2 receptor and cpEGFP. **(B)** The steps used to screen and optimize candidate PGE2 sensors, resulting in PGE2-1.0, as well as the development of a ligand-insensitive PGE2mut to serve as a negative control. **(C)** PGE2-1.0 and PGE2mut were expressed in HEK293T cells, and the change in fluorescence was measured in response to 10 μM PGE2. Scale bars, 20 μm. **(D)** Dose-response curves for PGE2-1.0 and PGE2mut expressed in HEK293T cells. **(E)** The left inset shows the structures of PGE2, PGD2, PGF2α, and PGI2, while the right inset shows the dose-response curves for PGE2-1.0 in response to these four prostaglandins. The main graph shows the normalized change in fluorescence in response to the indicated compounds (each applied at 1 μM, except the EP2 antagonist PF04418948, which was applied at 10 μM); n = 3 coverslips/group. **(F)** The 1-photon and 2-photon spectra of the PGE2-1.0 sensor measured in the absence (dashed lines) or presence (solid lines) of 10 μM PGE2. **(G)** The local puffing system, in which a glass pipette containing 10 μM PGE2 was positioned above an PGE2-1.0-expressing cell. The dotted black line indicates the ROI for line-scanning. Scale bar, 20 μm. **(H)** Left: a representative trace of the change in PGE2-1.0 fluorescence measured using line-scanning upon application of 10 μM PGE2 (top) and 100 μM EP2 antagonist (bottom); the solid green lines indicate the exponential fit to the data. Right: summary of the on (n = 10 cells/3 coverslips) and off (n = 6 cells/3 coverslips) time constants (τ_on and_ τ_off_, respectively). **(I)** Luciferase complementation assay for assessing downstream coupling to Gs proteins. Ctrl, Gs-LgBit alone; n = 3 cultures each. **(J)** Tango assay for assessing downstream coupling to β-arrestin. Ctrl, no receptor; n = 3 cultures each.

When expressed in HEK293T cells, PGE2-1.0 traffics to the plasma membrane and produces a dose-dependent increase in fluorescence in response to extracellular PGE2 application. In addition, the sensor’s mean (±SEM) half-maximal effective concentration (EC_50_) for PGE2 is 42±10 nM, similar to the EC_50_ reported for the wild-type EP2 receptor^11^. In contrast, as expected PGE2mut traffics to the plasma membrane, but has no detectable response to PGE2 application, even at 10 μM (Figure 1C, D). With respect to ligand specificity, the PGE2-1.0 sensor has a considerably lower EC_50_ (i.e., higher affinity) for PGE2 compared to the structurally similar prostaglandins PGD2, PGF2α, and PGI2 (with EC_50_ values of 24 μM, 195 μM, and 1 mM, respectively) (Figure 1E). In addition, the PGE2-induced response was fully blocked by the EP2 receptor antagonist PF04418948, and no detectable response was induced by any other lipid molecules tested such as docosahexaenoic acid (DHA) and sphingosine-1-phosphate (S1P), or by any classic neurotransmitters or neuromodulators tested (Figure 1E).

We also examined the sensor’s spectral properties, kinetics, and possible downstream coupling in HEK293T cells. Using one-photon excitation, the excitation peak of PGE2-1.0 is ∼500 nm with an isosbestic point at ∼420 nm, and the emission peak is 520 nm; using two-photon excitation, the excitation peak is ∼910 nm (Figure 1F). Using local perfusion and high-speed line-scanning (Figure 1G), we measured rapid response kinetics, with an average on-rate (τ_on_) of 0.4 s and an average off-rate (τ_off_) of 2 s (Figure 1H). To detect whether the PGE2-1.0 sensor engages downstream signaling pathways, we used the luciferase complementary assay and the Tango assay to measure activation of the GPCR-mediated Gs and β-arrestin pathways, respectively. As expected, the wild-type EP2 receptor couples robustly to both pathways; in contrast, PGE2-1.0 has only negligible downstream coupling in response to PGE2 (Figure 1I, J), confirming that expressing and activating the sensor does not affect the cell’s intrinsic signaling pathways.

To examine the performance of PGE2-1.0 when expressed in a physiologically relevant cell type, we expressed the sensor and the ligand-insensitive PGE2mut in cultured rat cortical neurons by infection with adeno-associated virus (AAV). We found that both PGE2-1.0 and PGE2mut are expressed widely throughout the plasma membrane, both on the cell body and on neurites (Figure 2A); moreover, the PGE2-1.0 sensor had a robust dose-dependent fluorescence response to PGE2 application, with an EC_50_ of 66±4 nM (Figure 2B, C). In contrast, PGE2mut produced no detectable response to PGE2 even at the highest concentration tested (Figure 2C). In addition, application of a saturating concentration of PGE2 led to a peak increase in PGE2-1.0 fluorescence of 1400%, with no measurable response in neurons expressing PGE2mut (Figure 2D). Similar to our results obtained with HEK293T cells, PGE2-1.0 had high specificity for PGE2 when expressed in neurons (Figure 2E). Finally, PGE2-1.0 produced a highly stable fluorescence response for up to 2 hours in the presence of 10 μM PGE2, indicating minimal sensor internalization or desensitization (Figure 2F–H). Overall, these results confirm that our PGE2-1.0 sensor can be used to reliably measure PGE2 dynamics *in vitro* with high sensitivity, specificity, and spatiotemporal resolution.

**Figure 2.**
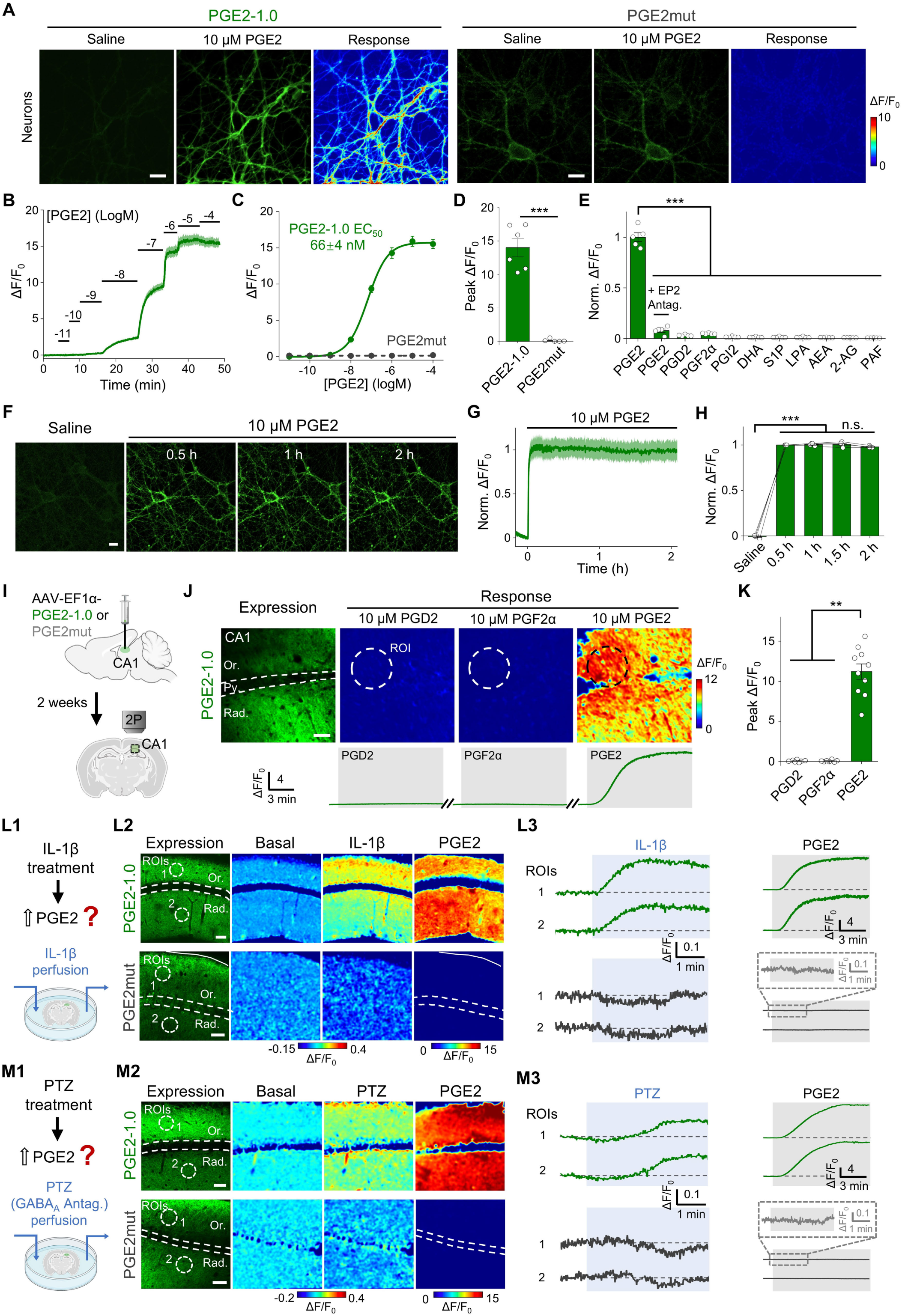
Characterization of PGE2-1.0 expressed in cultured neurons and acute brain slices. **(A)** PGE2-1.0 and PGE2mut were expressed in primary cultured cortical neurons, and the change in fluorescence was measured in response to 10 μM PGE2. Scale bars, 20 μm. **(B)** Example trace of the change in fluorescence in response to the indicated concentrations of PGE2 in primary cultured cortical neurons expressing PGE2-1.0; the data represent 30 ROIs measured in 1 coverslip. **(C)** Dose-response curve of PGE2-1.0 and PGE2mut in response to PGE2 measured in primary cultured cortical neurons. **(D)** Summary of peak ΔF/F_0_ measured in neurons expressing PGE2-1.0 (n = 6 coverslips) or PGE2mut (n = 5 coverslips). **(E)** Summary of the normalized change in PGE2-1.0 fluorescence measured in primary cultured cortical neurons in response to the indicated prostaglandins and lipids (each applied at 1 μM, except the EP2 antagonist, which was applied at 10 μM); n = 5 coverslips/group. **(F)** Example images of cultured neurons expressing PGE2-1.0, showing the fluorescence change in response to a 2-hour application of 10 μM PGE2; scale bar, 20 μm. **(G and H)** Example trace (G) and summary (H) of the normalized change in PGE2-1.0 fluorescence in the continued presence of 10 μM PGE2; n = 4 coverslips. **(I)** Schematic diagram depicting the strategy for injecting virus in the CA1 region of the mouse hippocampus, followed by fluorescence imaging of coronal slices using two-photon microscopy. **(J)** Top: expression of PGE2-1.0 in the CA1 region, and pseudo-color images of PGE2-1.0 in response to PGD2, PGF2α, and PGE2. Scale bar, 50 μm. The dashed circles show the ROIs used to measure the change in fluorescence, and the layers in the CA1 region are indicated (Or, stratum oriens; Py, stratum pyramidale; Rad, stratum radiatum). Bottom: traces of PGE2-1.0 fluorescence measured in the ROIs shown above, before and after application of PGD2, PGF2α, and PGE2. **(K)** Summary of the peak change in PGE2-1.0 fluorescence in response to PGD2, PGF2α, and PGE2 perfusion. **(L1)** Schematic drawing depicting the strategy for measuring the effect of IL-1β on PGE2 levels in brain slices expressing PGE2-1.0 or PGE2mut. **(L2)** Expression and pseudo-color images of the change in PGE2-1.0 and PGE2mut fluorescence in acute brain slices. The dashed circles indicate the ROIs used in L3, and the layers in the CA1 region are indicated. Scale bars, 50 μm. **(L3)** Example traces of the change in PGE2-1.0 and PGE2mut fluorescence in response to IL-1β (left) and PGE2 (right). **(M1)** Schematic drawing depicting the strategy for measuring the effect of GABA_A_ receptor antagonist pentylenetetrazol (PTZ) in brain slices expressing PGE2-1.0 or PGE2mut. **(M2)** Expression and pseudo-color images of the change in PGE2-1.0 and PGE2mut fluorescence in acute brain slices. Scale bars, 50 μm. **(M3)** Example traces of the change in PGE2-1.0 and PGE2mut fluorescence in response to PTZ (left) and PGE2 (right) application.

### PGE2-1.0 can be used to report endogenous PGE2 dynamics in acute brain slices

Next, we examined whether PGE2-1.0 can be used to monitor PGE2 dynamics in acute brain slices. We therefore injected AAV encoding PGE2-1.0 or PGE2mut in the hippocampal CA1 region of adult mice brain; after 2 weeks (to allow for expression), we prepared acute brain slices and measured fluorescence in the CA1 region using 2-photon microscopy (Figure 2I). We found that PGE2-1.0 responded specifically to PGE2, with an average peak response of ∼1100% when applied at 10 μM; in contrast, no response was measured for PGD2 or PGF2α (Figure 2J, K).

We then examined whether PGE2-1.0 can reliably detect changes in endogenous PGE2 in the hippocampal CA1 region. Given that PGE2 is closely related to inflammation^3,29^, we perfused the cytokine IL-1β on the slices to mimic an inflammation-like condition and measured the change in fluorescence (Figure 2L1). We found that IL-1β induced a significant increase in PGE2-1.0 fluorescence in both the stratum oriens and stratum radiatum layers of the CA1; in contrast, PGE2mut was expressed at high levels but had no measurable response to either IL-1β or PGE2 application (Figures 2L and S3). Similarly, given the close relationship between PGE2 synthesis and synaptic activity, including seizure activity^30^, we applied the GABA_A_ receptor antagonist pentylenetetrazol (PTZ) to mimic a seizure-like activity (Figure 2M1). We found that PTZ induced a significant increase in PGE2-1.0 fluorescence, while PGE2mut had no measurable response to PTZ or PGE2 (Figure 2M and S3). Taken together, these results confirm that our PGE2-1.0 sensor can reliably measure changes in endogenous PGE2.

### Detecting endogenous PGE2 dynamics *in vivo*

Based on our characterization of PGE2-1.0 in cultured cells and acute brain slices, we next asked whether this sensor can be used *in vivo* to detect endogenous PGE2 in the brain. PGE2 is known to mediate fever in the brain during inflammatory conditions^3,7^. Endogenous pyrogens and exogenous pyrogens such as lipopolysaccharide (LPS) can lead to an increase in PGE2 levels, which in turn can mediate hyperthermia or hypothermia (with high dose pyrogens) via EP3 receptors in the preoptic area (POA)^31–33^. Moreover, inhibiting PGE2 synthesis or knocking out the enzymes involved in its synthesis (primarily PTGES1 and COX-2) were shown to effectively reduce pyrogen-induced fever^19,32,34,35^. Therefore, to test the ability of PGE2-1.0 to measure changes in endogenous PGE2 in the brain in response to pyrogens, we performed fiber photometry recordings of PGE2-1.0 fluorescence while measuring body temperature in mice following LPS stimulation.

We first expressed PGE2-1.0 in the mouse brain by injecting the AAV into the medial preoptic area (MnPO), a subregion of the POA in the hypothalamus; we then implanted an optic fiber for subsequent recording (Figure 3A). We found that LPS (1 mg/kg body weight) induced a slow (over the course of 2–4 hours) but steady increase in PGE2-1.0 fluorescence, together with a decrease in the mouse’s skin temperature, indicating an increase in endogenous PGE2 levels in the MnPO (Figure 3B, C). To confirm the specificity of the PGE2-1.0 response, we pre-treated mice with the COX inhibitor indomethacin (Indo, 15 mg/kg body weight) 30 min prior to LPS administration and found that the LPS-induced PGE2-1.0 response and change in skin temperature were significantly reduced (Figure 3B, C). Thus, using this classic paradigm of LPS-induced acute inflammation, PGE2-1.0 has sufficient sensitivity and specificity to detect changes in endogenous PGE2 *in vivo*.

**Figure 3.**
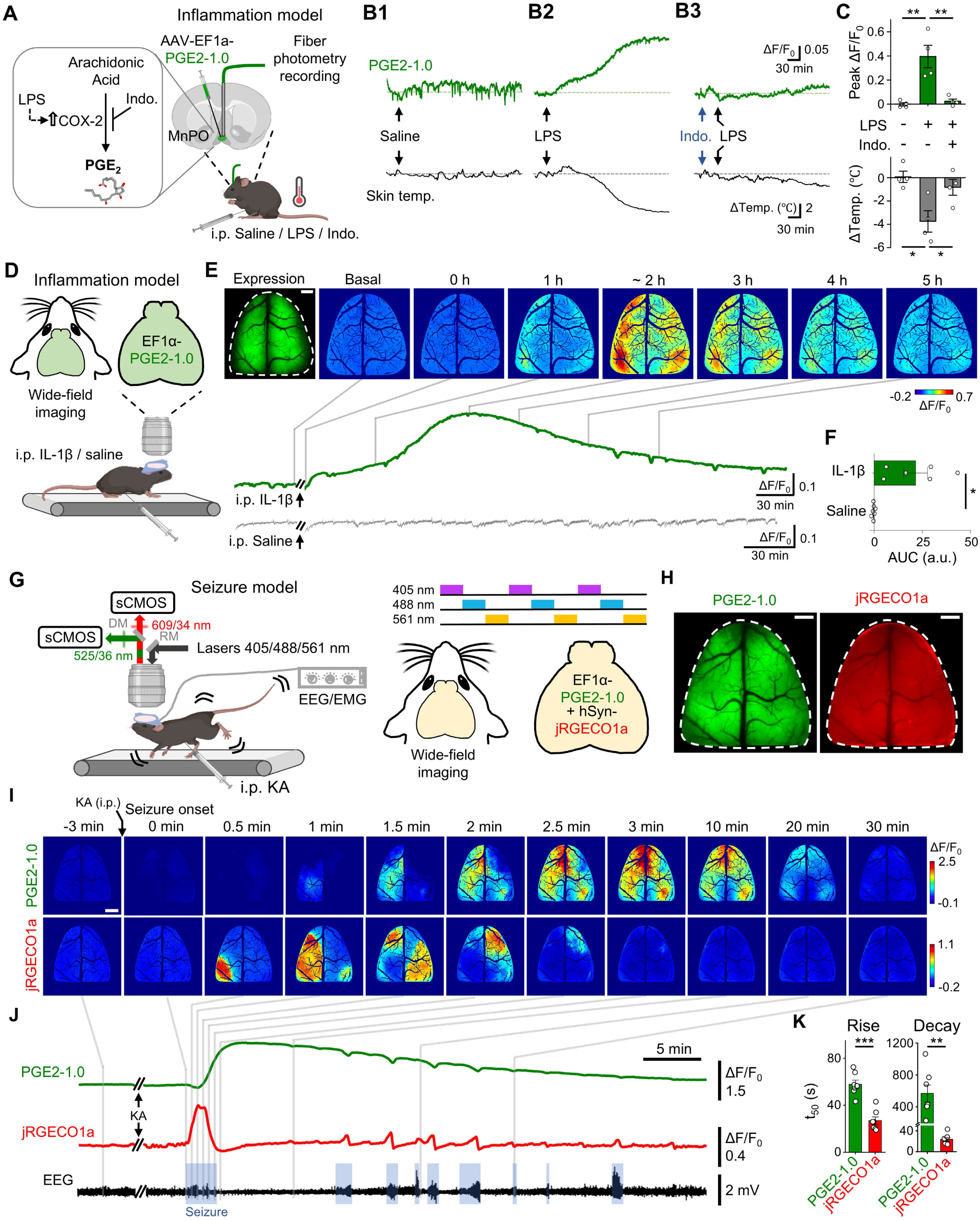
Using the PGE2-1.0 sensor to measure inflammation- and seizure-induced changes in PGE2 *in vivo*. **(A)** Left: diagram depicting the pathway for PGE2 synthesis and compounds that block or increase the activity of specific components in this pathway. LPS, lipopolysaccharide; COX-2, cyclooxygenase-2; Indo., indomethacin. Right: schematic diagram depicting the strategy for injecting virus, implanting optic fibers, and fiber photometry recording in the MnPO region of the mice brain; the animal’s skin temperature was also recorded during the experiment. **(B)** Example traces of the change in PGE2-1.0 fluorescence and skin temperature in response to an i.p. injection of saline (B1), 1 mg/kg body weight LPS (B2), or 1 mg/kg body weight LPS following a pre-injection of 15 mg/kg body weight indomethacin (B3). **(C)** Summary of the peak change in PGE2-1.0 fluorescence and skin temperature (n = 4 mice). **(D)** Schematic diagram depicting the strategy used for wide-field imaging; PGE2-1.0 was expressed throughout the cortical region, and the mice were head-fixed for wide-field imaging. **(E)** Top: expression and pseudo-color images of the change in PGE2-1.0 fluorescence at the indicated times; where indicated, IL-1β (25 μg/kg body weight) or saline was injected i.p. Scale bar, 1 mm. Bottom: corresponding example traces of PGE2-1.0 fluorescence. **(F)** Summary of the area under the curve (AUC, in arbitrary units) of the change in PGE2-1.0 fluorescence measured for 240 min after injecting IL-1β (n = 6 mice) or saline (n = 7 mice). **(G)** Schematic diagram depicting the strategy for dual-color wide-field imaging in a kainic acid (KA)-induced seizure model. sCMOS, scientific complementary metal oxide semiconductor camera; DM, dichroic mirror; RM, reflective mirror. **(H)** Expression of PGE2-1.0 and jRGECO1a in the mouse cortex. Scale bars, 1 mm. **(I)** Representative images of the change in PGE2-1.0 and jRGECO1a fluorescence in response to an i.p. injection of KA (10 mg/kg body weight). **(J)** Traces of the change in PGE2-1.0 and jRGECO1a fluorescence, together with the corresponding EEG signal. The blue shaded boxes in the EEG trace indicate epileptic discharges. **(K)** Summary of the rise and decay kinetics of the change in PGE2-1.0 and jRGECO1a fluorescence measured during seizure activity; n = 6 mice.

### Using PGE2-1.0 to monitor inflammation- and seizure-induced changes in cortical PGE2 with high spatiotemporal resolution

Although PGE2 and its receptors are widely distributed throughout the brain^8,10,9^, dynamic changes in PGE2 in specific brain regions are seldom studied due to the lack of suitable detection methods. Taking advantage of the PGE2-1.0 sensor’s high specificity and spatiotemporal resolution, we expressed the sensor in the mouse cerebral cortex and then used mesoscopic imaging to measure endogenous PGE2 during various paradigms such as inflammation and seizure activity.

We first expressed the PGE2-1.0 sensor homogeneously throughout the cerebral cortex and then measured the change in fluorescence in response to an i.p. injection of IL-1β (25 μg/kg body weight) or saline (Figure 3D). We found that IL-1β caused a gradual increase in PGE2-1.0 fluorescence that peaked at ∼22% above baseline and then slowly returned to baseline levels; in contrast, saline had no effect on PGE2-1.0 fluorescence (Figure 3E, F, Video S1).

Next, we examined PGE2 dynamics in a kainic acid (KA)-induced seizure model. PGE2 has long been studied for its close connection with the onset and progression of seizure activity^36–39^, but little is currently known regarding the spatiotemporal changes in PGE2 levels during seizure, much less the relationship between PGE2 and other seizure-related signals such as calcium (Ca^2+^). To address these questions, we co-expressed PGE2-1.0 and the red fluorescent Ca^2+^ sensor jRGECO1a in the cerebral cortex; in addition, we implanted electroencephalography (EEG) and electromyography (EMG) electrodes to monitor seizure activity (Figure 3G). After allowing the mice to recover for two weeks, we used a custom-built wide-field microscope to record both PGE2-1.0 and jRGECO1a fluorescence together with the EEG and EMG signals (Figure 3G). Dual-color imaging confirmed that both PGE2-1.0 and jRGECO1a were expressed homogeneously throughout the dorsal cortex (Figure 3H). After allowing the mouse to adapt to the wide-field imaging device, we gave an i.p. injection of KA (10 mg/kg body weight). During seizure activity, we measured a transient increase in Ca^2+^ that started in a relatively small initial area and then rapidly spread to surrounding regions throughout the entire cortex; in contrast, the PGE2 signal did not propagate, but increased gradually throughout the cortex over time after seizure onset (Figure 3I, Video S2), thus revealing a spatial pattern distinct from the Ca^2+^ signal. In addition, the PGE2 and Ca^2+^ signals had distinct temporal features— immediately after seizure onset, we observed a rapid rise and fall in the Ca^2+^ signal, while the PGE2-1.0 signal began slightly later than the Ca^2+^ signal, increased relatively slower, and reached a maximum response of ∼150% that was maintained for several minutes (Figure 3I, J). To quantify this difference, we measured the rise and decay kinetics (t_50_) of the PGE2 and Ca^2+^ signals and found that the PGE2 signal was significantly slower than the Ca^2+^ signal (Figure 3K). Thus, our results reveal significant differences in the spatiotemporal patterns of PGE2 and Ca^2+^ during seizure activity.

Interestingly, we also observed notable differences in the spatiotemporal dynamics of PGE2 between the inflammation and seizure models. First, in terms of the temporal dynamics, we found that both the rise and decay kinetics of PGE2 were significantly faster in the seizure model compared to the inflammation model (Figure 4A, B). Second, the average peak change in PGE2 during seizure was significantly larger compared to inflammation (Figure 4A, C). It is worth noting that although PGE2 has long been studied for its close connection with the onset and progression of seizure, it was previously considered to participate primarily in the chronic inflammation that accompanies seizure^36–39^. Thus, unlike traditional methods used to monitor PGE2, which are unable to capture rapid changes, the high temporal resolution of our PGE2-1.0 sensor makes it a robust new tool for detecting rapid changes in endogenous PGE2 levels.

**Figure 4.**
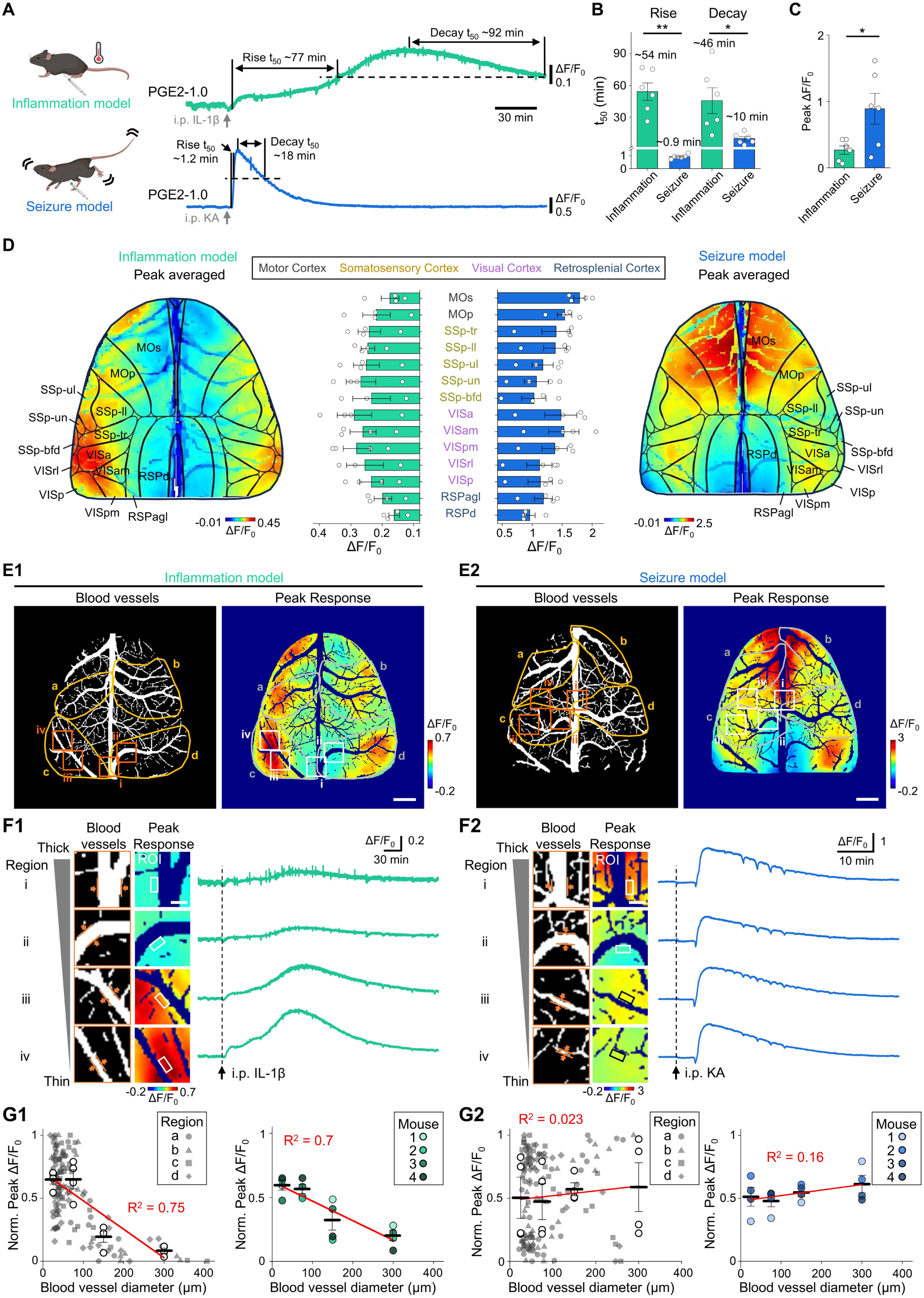
The PGE2-1.0 sensor reveals distinct temporal and spatial differences in PGE2 dynamics in the brain. **(A)** Schematic diagram depicting the IL-1β-induced inflammation and KA-induced seizure models (left), and representative traces of the change in PGE2-1.0 fluorescence in response to an i.p. injection of IL-1β or KA (right). In the traces, the rise and decay t_50_ values are indicated. **(B)** Summary of the rise and decay t_50_ values for the change in PGE2 fluorescence measured as shown in (A); note the break in the *y*-axis; n = 6 mice. **(C)** Summary of the peak change in PGE2 fluorescence measured during inflammation and seizure; n = 6 mice. **(D)** Pseudo-color images depicting the average peak increase in PGE2-1.0 fluorescence measured during inflammation (left) and seizure (right); shown in the middle are the peak responses measured in the indicated cortical structures; n = 4 mice. **(E)** Representative images of the cortical blood vessel structures (left) and the peak increase in PGE2-1.0 fluorescence (right) measured during inflammation (E1) and seizure (E2); the ROIs used in (F) and (G) are indicated. Scale bars, 1 mm. **(F)** Magnified views of the images of region i–iv shown in (E) (left) and traces of the change in PGE2 fluorescence measured at the ROIs indicated by the rectangles (right). Scale bar, 200 μm. **(G)** Left: normalized peak increase in PGE2-1.0 fluorescence plotted against the adjacent blood vessel diameter in the ROIs shown in (E) in the inflammation (G1) and seizure (G2) models; also shown are the correlation coefficients. Right: normalized peak increase in PGE2-1.0 fluorescence plotted against the adjacent blood vessel diameter for 4 mice; also shown are the correlation coefficients.

With respect to the spatial patterns, we also found key differences between the two disease models, with notable differences in the location of the resulting signals; specifically, the seizure-induced changes in PGE2 levels were largely observed in the anterior cortex, while the inflammation-induced changes in PGE2 did not appear to differ among various subcortical regions (Figure 4D). In the acute inflammation model, the area with the largest response was relatively close to the peripheral regions of the cortex, near the ends or bifurcations of large blood vessels (Figure 4E1); in contrast, the area with the largest response in the seizure model was closer to anterior brain regions (Figure 4E2). An analysis of the correlation between the PGE2-1.0 response and the diameter of adjacent blood vessels revealed that the inflammation-induced change in PGE2 was relatively larger near small-diameter vessels (Figure 4F1, G1); in contrast, we found no correlation between the seizure-induced change in PGE2 and the diameter of adjacent blood vessels (Figure 4F2, G2). Together, these results indicate that our new PGE2-1.0 sensor has sufficient temporal and spatial resolution to reveal highly detailed information regarding the dynamic changes in PGE2 under physiological and pathological conditions.

## DISCUSSION

Here we report the development and characterization of a genetically encoded fluorescent sensor for detecting PGE2 with high specificity, rapid kinetics, and high spatiotemporal resolution. We also show that this sensor can be used to reliably monitor changes in endogenous PGE2 both *in vitro* and *in vivo*.

Using fiber photometry and mesoscopic imaging, we observed that PGE2 levels increase relatively slowly following an i.p. injection of the pro-inflammatory agents LPS and IL-1β, reaching peak levels after 2–4 hours, consistent with previous studies^31^. This finding may suggest that specific brain regions rely on similar—or closely related—mechanisms for the production and release of PGE2 in response to acute inflammation. In addition, we measured PGE2 during seizure activity and found a rapid increase (within ∼2 min) in extracellular PGE2 levels following the onset of seizure. Previous studies focused primarily on the long-term increase in PGE2, for example 24 hours or several days after seizure, and examined the associated chronic inflammation^30,40^. However, our results indicate that robust changes in PGE2 levels begin much earlier after seizure onset, providing future directions for the study of seizure mechanisms and treatments. The major differences in the timing of PGE2 release between the acute inflammation and seizure models may also suggest differences in the mechanisms underlying the production and/or release of PGE2 in these two distinct disease states.

With respect to the spatial changes in PGE2, we found notable differences in cortical PGE2 changes between the inflammation and seizure models. Specifically, acute inflammation induced a relatively large response close to the cortical periphery, near major blood vessel endings or bifurcations, while seizure induced the largest response near anterior brain regions. Previous studies suggest that under acute inflammatory conditions, LPS or IL-1β may act either directly or indirectly on vascular endothelial cells in the brain to induce PGE2 synthesis, while knocking out COX-2 or the terminal enzyme for PGE2 synthesis in endothelial cells can prevent LPS-induced fever^41,42,35^. However, none of these previous studies measured PGE2 dynamics directly; rather, they used body temperature and/or COX-2/PTGES expression as an indirect proxy for measuring PGE2 dynamics. Our results obtained using PGE2-1.0 fluorescence imaging may therefore provide evidence to support the hypothesis that PGE2 originates in the vascular endothelium.

In summary, our genetically encoded PGE2-1.0 sensor can be used to reliably measure changes in endogenous PGE2 with high specificity, sensitivity, and spatiotemporal resolution, providing a robust new tool for studying PGE2 dynamics both *in vitro* and *in vivo*.

## ACKNOWLEDGMENTS

This research was supported by grants from the National Natural Science Foundation of China (31925017), the National Key R&D Program of China (2022YFE0108700 and 2023YFE0207100), Beijing Municipal Science & Technology Commission (Z220009); grants from the NIH BRAIN Initiative (1U01NS120824), the Feng Foundation of Biomedical Research, the Clement and Xinxin Foundation, the New Cornerstone Science Foundation through the New Cornerstone Investigator Program (to Y.L.); and grants from the Peking-Tsinghua Center for Life Sciences and the State Key Laboratory of Membrane Biology at Peking University School of Life Sciences (to Y.L.).

We thank Xiaoguang Lei at PKU-CLS and the Optical Imaging platform and small animal imaging platform of National Center for Protein Sciences at Peking University in Beijing, China, for their support and assistance with the Opera Phenix, the Operetta CLS high-content imaging system, the Nikon A1RSi+ laser scanning microscope, and the behavior facility. We thank the Laboratory Animal Center of Peking University for advice and technical support.

## AUTHOR CONTRIBUTIONS

Y.L. supervised the study. L.W. and Y.Y. performed the experiments related to sensor development, optimization, characterization in HEK293T cells and cultured neurons, sensor validation on acute brain slices, and *in vivo* fiber photometry as well as wide-field imaging experiments in living mice. Y.Q. performed characterization of the sensor’s downstream coupling. F.D. initiated and helped with experiments related to *in vivo* wide-field imaging. All authors contributed to the interpretation and analysis of the data. L.W. and Y.L. wrote the manuscript with contributions from all authors.

## DECLARATION OF INTERESTS

All authors declare no competing interests.

## SUPPLEMENTAL INFORMATION

**Document S1**. Figures S1–S3

**Video S1**. Cortex-wide imaging of IL-1β-induced PGE2 dynamics in vivo, related to Figure 3

**Video S2**. PGE2 and calcium signal dynamics during seizure, related to Figure 3

## METHODS

### Cell lines

HEK293T cells were cultured in DPF medium (high-glucose DMEM supplemented with 10% fetal bovine serum, 100 U/mL penicillin, and 0.1 mg/mL streptomycin) at 37°C in humidified air containing 5% CO_2_. Cells at 60–70% confluence were transfected with a 3:1 mass ratio of polyethyleneimine (PEI) to plasmid DNA. In brief, PEI and plasmid DNA were combined in a microtube, and after 15 min at room temperature the PEI-DNA mixture was added to the cells; the medium was changed approximately 6 hours later. The cells were then allowed to express the plasmid DNA for 24–48 hours before imaging.

### Neuronal cultures and viral infection

Primary neurons were derived from P0 Sprague-Dawley rat pups. The cerebral cortex was dissected and cortical neurons were dissociated by using 0.25% trypsin-EDTA. Then, neurons were plated on 12 mm diameter glass coverslips (previously sterilized and coated with poly-D-lysine) and cultured in Neurobasal medium supplemented with 2% B-27, 1% GlutaMAX, 100 U/mL penicillin, and 0.1 mg/ml streptomycin; every 3 days, half of the medium was replaced with fresh medium. Adeno-associated virus (AAV, 1 μl, at a titer of about 3×10^12^ vg/ml) was added to the neurons at 3–7 days *in vitro* (DIV3–7), and imaging was performed on DIV13–25.

### Measurements of spectra

For one-photon spectra, HEK293T cells expressing PGE2-1.0 was collected and transferred to a 384-well plate in the absence or presence of 10 μM PGE2. Excitation and emission spectra were measured at 5 nm increments with a 20 nm bandwidth using a Safire2 multi-mode plate reader (Tecan). Control cells not expressing a sensor were prepared to the same density and were measured using the same protocol for background subtraction. For two-photon spectra, HEK293T cells expressing PGE2-1.0 were cultured on 12 mm coverslips. The two-photon excitation spectra was measured at 10 nm increments and ranging from 700 to 1050 nm using an Ultima Investigator two-photon microscope (Bruker) equipped with a × 20/1.0 NA water-immersion objective (Olympus), an InSight X3 tunable laser (Spectra-Physics) and the Prairie View 5.5 software (Bruker).

### High-content imaging platform

HEK293T cells cultured in CellCarrier Ultra-Black 96-well plates were imaged using an Opera Phenix high-content imaging analysis system (PerkinElmer) equipped with 20× air, 40× air, and 40× water-immersion objectives, using 488 nm and 561 nm lasers for excitation. The signals from the green and red fluorescent proteins were collected via 525/50-nm and 600/30-nm filters, respectively. The accompanying software package Harmony (PerkinElmer) was used to analyze the fluorescence signals, localizing the cell membranes in the field of view using the membrane-targeted mCherry red fluorescence signal and delineating regions of interest (ROIs). The ratio between green and red fluorescence intensity was calculated and used to measure the brightness of the green sensor (i.e., the corrected F value). The change in the sensor’s fluorescence intensity using the formula ΔF/F_0_, where F_0_ is baseline fluorescence.

### Luciferase complementation assay

The HEK293T cells were transfected with the wild-type EP2 receptor, the PGE2-1.0 sensor, or Gs-LgBit alone. The luciferase complementation assay was performed as previously described^43^. In brief, 24–36 h after transfection, the cells were dissociated using a cell scraper, resuspended in PBS, and transferred to 96-well plates. Then, 5 μM furimazine (NanoLuc Luciferase Assay, Promega) and PGE2 at various concentrations (ranging from 0.01 nM to 10 μM) were bath-applied to the cells. After incubation for 10 min in the dark, luminescence was measured using a VICTOR X5 multilabel plate reader (PerkinElmer).

### Tango assay

A reporter cell line, HTLA cells that stably expressing a tTA-dependent luciferase reporter and a β-arrestin2-TEV fusion gene, was transfected with pTango vectors to express wild-type EP2 receptor or the PGE2-1.0 sensor, and the empty vector was used as a control. At 24 h after transfection, the cells were transferred to 96-well plates and bathed with PGE2 at varying concentrations (ranging from 0.01 nM to 30 μM). The cells were then cultured for 12 h to allow the expression of tTA-dependent luciferase. Bright-Glo reagent (Fluc Luciferase Assay System, Promega) was added to a final concentration of 5 μM, and luminescence was measured using a VICTOR X5 multilabel plate reader (PerkinElmer).

### Confocal microscopy

Confocal microscopy was performed using a Nikon Ti-E A1 laser-scanning confocal microscope equipped with a 10×/0.45-NA objective, a 20×/0.75-NA objective, and a 40×/1.35-NA oil-immersion objective; a 488 nm laser was used for excitation, and fluorescence was collected via a 525/50-nm filter. During imaging, the coverslip was placed in a custom-made imaging chamber at room temperature (24°C). The microscope was controlled using NIS-Elements (Nikon), and the data were processed and analyzed using ImageJ software.

### Animals

All animal protocols were approved by the respective laboratory animal care and use committees at Peking University. The mice used were both male and female C57BL/6J mice (Charles River Laboratories). For these experiments, P0 and adult (age 6–12 months) mice were used; these mice were housed in a temperature-controlled animal facility with a 12-hour light/dark cycle, with food and water available ad libitum. P0 Sprague-Dawley rats (Charles River Laboratories) of both sexes were used to prepare primary neuronal cultures as described above.

### Stereotaxic AAV injection and optic fiber implantation

To anesthetize adult mice, 2,2,2-tribromoethanol (Avertin, 240 mg/kg body weight, Sigma-Aldrich) was administered by i.p. injection. The anesthetized mice were then secured in a stereotaxic frame for AAV injection. Using a microinjection pump, 300 nL of virus (AAV2/9-EF1α-PGE2-1.0, 2.68×10^12^ vg/ml, packaged by BrainVTA) was slowly injected into the median preoptic nucleus (MnPO) of the hypothalamus using the following coordinates: AP, 0 mm relative to Bregma; ML, -0.6 mm relative to Bregma; and DV, -5 mm. After AAV injection, the needle was left in place for 5 min, and an optic fiber was implanted at the injection site. The entire setup was secured with dental cement, and recordings were performed after 14 days to allow for viral expression.

### Fiber photometry recording

To collect the signal from the PGE2-1.0 sensor, we used a FPS-410/470/561 photometry system (Inper) with the Inper signal v.2.0.0 software (Inper). In brief, a 10-Hz (with 20-ms pulse duration) 470/5-nm filtered light-emitting diode (LED) at 20 μW was used to excite the green fluorescent and the emission light were collected at 520 nm. For recording, the mouse was connected to the fiber optic system and allowed to freely move around and acclimate in a behavioral chamber for approximately 30 min, which the recorded fiber optic signals served as the baseline.

In the experiment inducing endogenous PGE2 release with lipopolysaccharide (LPS), a stock solution of LPS (5 mg/mL; Sigma, L4130) was diluted with saline and injected i.p. at 1 mg/kg body weight. For the cyclooxygenase inhibitor indomethacin (Indo, MedChemExpress, 53-86-1) the stock solution (10 mM) was diluted with 0.2 M Tris-HCl (pH 8) and injected i.p. at 15 mg/kg body weight. For IL-1β (Novoprotein, C042) the stock solution (100 μg/mL) was diluted with saline and injected i.p. at 25 μg/kg body weight.

### Mice body temperature recording

During experiments, the mice were placed in the behavioral chamber situated in a temperature-controlled room, with an infrared thermal camera (Inf iRay, AT20) fixed at the top of the behavsioral chamber to monitor mice skin temperature through infrared imaging. The field of view of this camera covers the entire behavioral chamber so the free-moving mice could be monitored at any position. This camera could continuously monitor the temperature within its field of view and generate temperature readings at one-minute intervals. Each reading included the maximum, minimum, and average temperatures observed within the field of view. The maximum temperature recorded at each time point was utilized as an indication of the mice’s skin temperature.

### Wide-field imaging surgeries

Newborn (P0–P1) C57BL/6J pups received an injection of a 1:1 virus mixture containing AAV2/9-EF1α-PGE2-1.0 (2.68×10^12^ vg/ml, packaged by BrainVTA) and AAV2/9-hSyn-NES-jRGECO1a (8.3×10^13^ vg/ml, packaged by WZ Biosciences Inc.) in the transverse sinus on both hemispheres (3 μl per hemisphere) via a microinjection pump (KD Scientific, LEGATO 130). Prior to virus injection, the pups were anesthetized by placing them on ice for 1–2 min. When the pups stopped moving, they were then placed on a cold metal plate with ice packs to keep them cool, and their heads were secured. The skin on both sides of the transverse sinus was incised, and a glass electrode was used to puncture the wall of the transverse sinus, ensuring that the electrode tip was located inside the sinus. The virus mixture was then injected at a rate of 1.2 μl/min, after which the skin at the injection site was closed using tissue adhesive (3M Vetbond, 1469SB). The pups were then placed on a warming pad and monitored until they recovered; when they resumed normal activity, they were returned to their home cage.

The craniotomy procedure and implantation of EEG and EMG electrodes were performed 8–10 weeks after virus injection. The mice were anesthetized with Avertin (240 mg/kg body weight) and secured in a stereotaxic injector while maintaining inhalation anesthesia with 1% isoflurane. Erythromycin ointment was applied to the eyes, and the fur and scalp were removed from the top of the head, followed by cleaning the underlying connective tissue. A cranial drill was used to create an 8 mm × 8 mm window in the skull, and care was taken to remove the skull section. A custom-made thin glass coverslip of the same shape as the cranial window was affixed in place to replace the skull section and serve as an imaging window. Four EEG electrodes were also implanted: two near the frontal cortex in each hemisphere, one near the right hippocampal cortex, and one near the cerebellum. Finally, two EMG electrodes were inserted into the neck muscle tissue. The electrode micro-connectors were glued to the skull using cyanoacrylate adhesive, and a custom metal headpiece was attached to the mouse’s head for subsequent head stabilization. The entire setup was secured with dental cement, while ensuring that the craniotomy window was not obscured. The mice were allowed to recover for 1–2 weeks, after which wide-field imaging experiments were performed.

A custom-built two-color wide-field microscope equipped with a 2×/0.5-NA objective (Olympus, MVPLAPO2XC) and two 1×/0.25-NA tube lenses (Olympus, MVPPLAPO1X) was used to expand the field of view. The signal was acquired using two high-speed, high-sensitivity sCMOS cameras (Andor, Zyla 4.2 PLUS). The excitation light was delivered via a fiber-coupled laser (Changchun New Industries Optoelectronics Technology Co., Ltd.) capable of emitting excitation light at 405 nm, 488 nm, and 561 nm. The emission light was passed through a 567 nm long-pass dichroic beam splitter (Thorlabs, DMLP567L) for spectral separation, and then through either a 525/36 nm or 609/34 nm emission filter (Chroma) before entering the green and red channels, respectively, of the sCMOS cameras. Images were acquired using Micro-Manager 2.0 software with a resolution of 512×512 pixels, 4×4 pixel binning, and an exposure time of 40 msec. The laser and imaging system were coordinated and triggered using a script programmed into an Arduino board (Arduino Uno R3). During imaging, the mouse’s head was fixed to the base using the metal headpiece, and the EEG and EMG electrodes were connected, while a treadmill allowed the mouse to run freely. For the inflammation and seizure experiments, baseline data were recorded for ∼30–60 min, followed by an i.p. injection of either IL-1β (25 μg/kg body weight, to induce inflammation) or kainic acid (10 mg/kg body weight, to induce seizure); data were then recorded for 6 hours. Data from the wide-field imaging, EEG, EMG, running behavior, and infrared video recordings were synchronized using a Power1401 interface (Cambridge Electronic Design) for subsequent analysis.

### Data processing and statistics

In this study, data were analyzed and plotted using Microsoft Excel, Origin Pro, GraphPad Prism, Spike2, or MATLAB. Except where indicated otherwise, all summary data are presented as the mean ± the standard error of the mean (SEM). Data were analyzed using Student’s *t*-test or one-way ANOVA. Where indicated, \**p* < 0.05, \*\**p* < 0.01, \*\*\**p* < 0.001, and n.s., not significant (*p* > 0.05).

## Notes

### Competing Interest Statement

The authors have declared no competing interest.

