## Supplementary Figures for "A genetically encoded fluorescent sensor for monitoring spatiotemporal prostaglandin E2 dynamics *in vivo*"

### SUPPLEMENTARY FIGURE LEGENDS

**Figure S1. Screening of the $GRAB_{PGE2}$ sensor GPCR scaffold and insertion sites.** **(A)** Representative images of HEK293T cells co-expressing red fluorescent RFP-CAAX (to label the plasma membrane) and prototype $GRAB_{PGE2}$ sensor candidates with cpEGFP inserted in the indicated PGE2 receptor backbones. The dotted lines in the merged images were used to measure the correlation between the green and red fluorescence signals. Scale bar, 20 μm. **(B)** Line-scan traces of the green (cpEGFP) and red (RFP-CAAX) channels in the images shown in (A). **(C)** Summary of the correlation between the green and red fluorescence signals in order to confirm membrane trafficking of the various $GRAB_{PGE2}$ sensor candidates. **(D)** Peak $\Delta F/F_0$ plotted against maximum brightness for the indicated $GRAB_{PGE2}$ candidates. The candidate with the highest performance (PGE2-0.1) is indicated which was the selected candidate for further optimization.

**Figure S2. The structure and mutation sites in the PGE2-1.0 sensor.** **(A)** Structure of the PGE2-1.0 sensor, with the PE2 GPCR and cpEGFP components shown in purple and green, respectively. The linker and cpEGFP regions in which mutations were introduced to generate PGE2-1.0 are indicated. **(B)** The amino acid sequence of the PGE2-1.0 sensor, with the sensor's domains shown above. The mutation sites for generating the sensor are indicated by red boxes, and the $T113^{2.54}W$ mutation introduced to create the ligand-insensitive PGE2mut version from PGE2-1.0 is indicated by a black box. Note that the amino acid numbering system used here corresponds to the start of the IgK leader sequence.

**Figure S3. Summary of the change in PGE2-1.0 and PGE2mut fluorescence measured in brain slices in response to IL-1β, PTZ, and PGE2 perfusion.** **(A)** Summary of the peak change in PGE2-1.0 and PGE2mut fluorescence in brain slices in response to IL-1β, PTZ, and PGE2 perfusion. **(B)** Summary of the area under the curve (AUC) of the change in PGE2-1.0 and PGE2mut fluorescence measured 4 min after perfusing IL-1β or PTZ, and 10 min after perfusing PGE2.

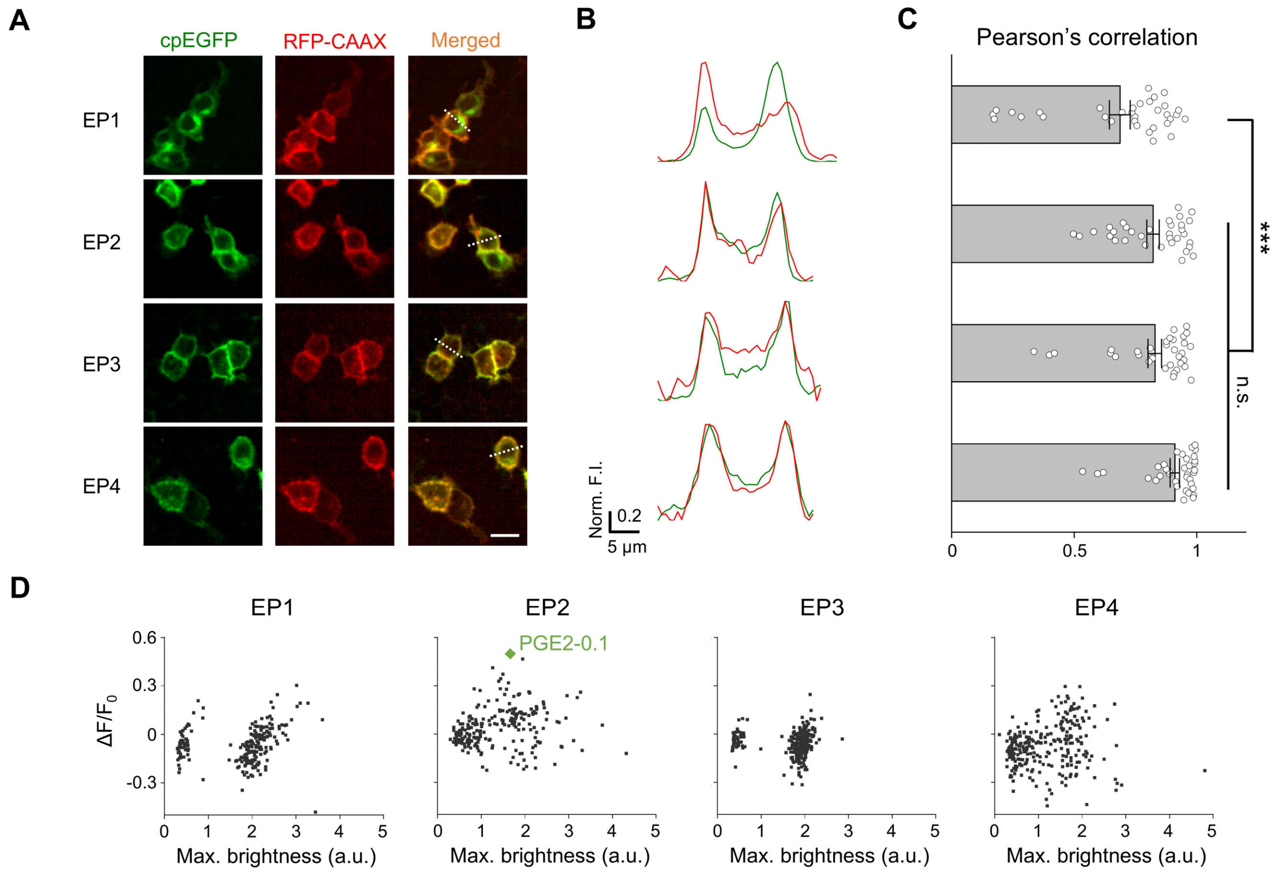


**Figure S1. Screening of the GRAB$_{PGE2}$ sensor GPCR scaffold and insertion sites.**

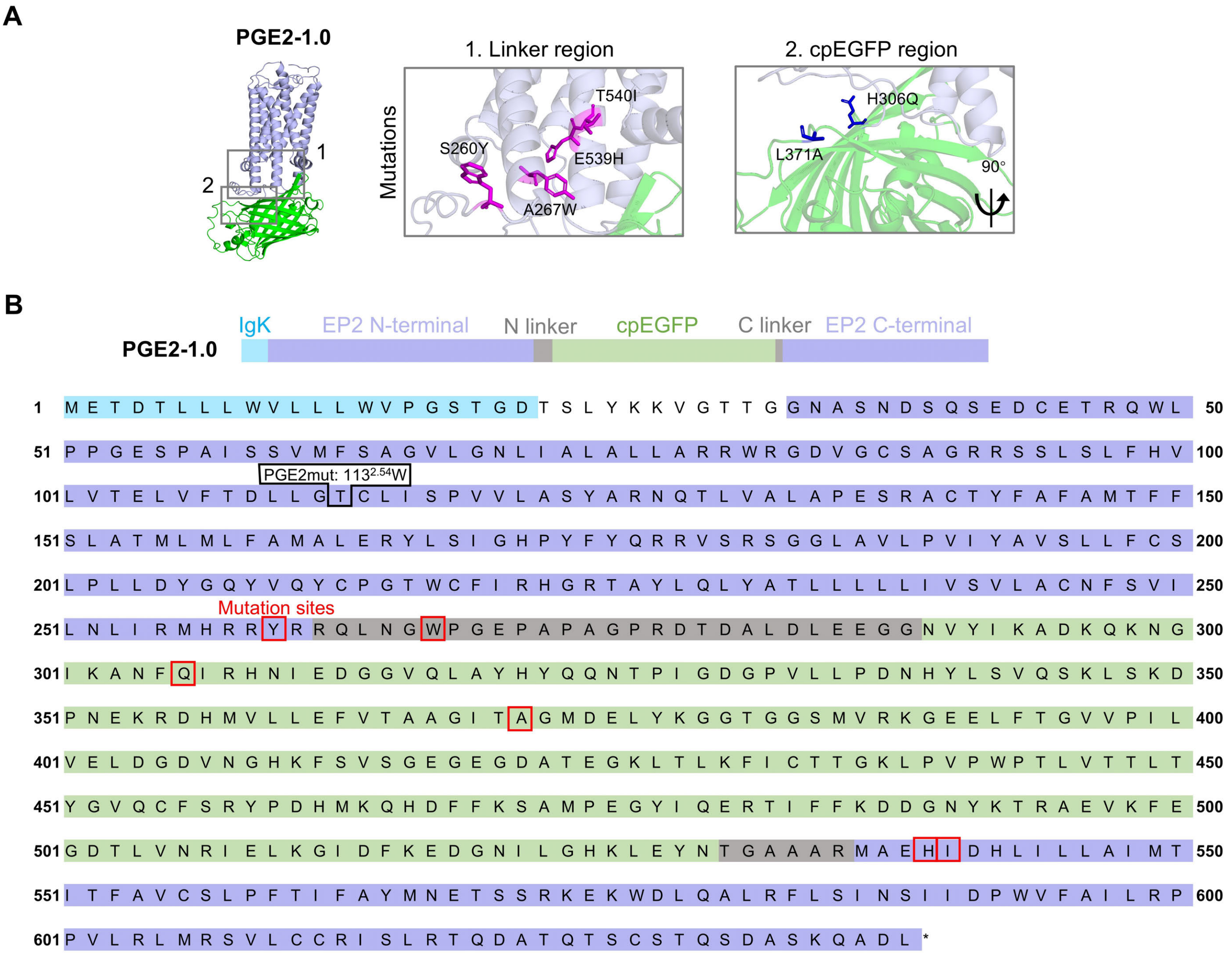


**Figure S2. The structure and mutation sites in the PGE2-1.0 sensor.**

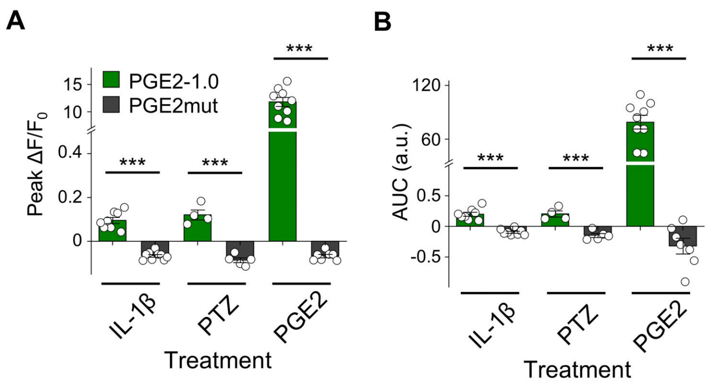


**Figure S3. Summary of the change in PGE2-1.0 and PGE2mut fluorescence measured in brain slices in response to IL-1β, PTZ, and PGE2 perfusion.**
